# A Systems Theory of Cardiac-Autonomic Trajectories and Equilibria

**DOI:** 10.64898/2026.11.02.763675

**Authors:** Matías Castillo-Aguilar, Diego Mabe-Castro, Cristian Núñez-Espinosa

## Abstract

Heart rate variability (HRV) is conventionally summarized with static indices computed over fixed windows, which cannot show how cardiac autonomic regulation persists, transitions, responds to perturbation, and recovers. We develop the Systems Theory of Cardiac-Autonomic Trajectories and Equilibria (STATE), in which autonomic regulation is a low-dimensional latent stochastic dynamical system observed only through RR intervals, and we derive what the theory can and cannot claim. First, we prove that for isotropic linear dynamics the sympathetic and parasympathetic coordinates are not separately identifiable from a scalar RR signal, whereas distinct time scales or coupling restore observability. Second, we derive exact properties of a minimal bistable system (stationary law, mean escape time, critical forcing), and a Monte Carlo simulation agreed with the exact escape time within 6%. Third, in 2000 simulated replicates of a monostable and a bistable system that share the same linearization at rest, baseline variability (area under the receiver operating characteristic curve, AUC, 0.56) and peak displacement at the pulse used (AUC 0.42) did not discriminate the systems, yet sustained return after an identical pulse differed: 100% versus 7%. We define resilience as a multidimensional response profile, show that recovery descriptors depend on analyst-chosen tolerances that must be fixed a priori, and state four falsifiable hypotheses with the result that would refute each. The work is theoretical and computational; no empirical recordings were used.

## 1 Introduction

At the organ level, cardiac rhythm is generated by pacemaking cells in the sinoatrial (SA) node, which continuously integrate competing and synergistic neural signals from sympathetic postganglionic fibers (sympathetic nervous system, SNS) and parasympathetic vagal fibers (parasympathetic nervous system, PNS). This peripheral interface is regulated by subcortical and cortical structures within the central autonomic network (Benarroch 1993), coupled with arterial baroreflex and respiratory feedback loops (Eckberg 1983). Heart rate variability (HRV) therefore reflects the output of a continuous, nonlinear neurocardiac regulatory process operating across multiple time scales.

Despite this dynamic complexity, cardiac autonomic function is conventionally quantified through summary statistics (such as SDNN, RMSSD, or low- and high-frequency spectral power) (Task Force of the European Society of Cardiology and the North American Society of Pacing and Electrophysiology 1996; Shaffer and Ginsberg 2017) computed over fixed windows and treated as approximately stationary within them. Recent reviews underscore that, unlike static biomarkers, HRV is highly sensitive to measurement context, recording duration, and physiological state, so isolated summary values cannot be read as context-free indicators of autonomic health (Ghazzal et al. 2026; Sundas et al. 2025). These indices are useful, but they collapse a continuously evolving process into point estimates and lose information on how the system moves through state space: whether autonomic modulation settles into one or several stable regimes, how far an internal or external perturbation displaces it, and how rapidly it recovers. Two individuals with identical RMSSD or mean RR over a 5-minute recording may nonetheless have fundamentally different regulatory architectures, perturbation sensitivities, and recovery trajectories.

An alternative is to treat cardiac autonomic regulation explicitly as a latent dynamical system: a time-varying state, observed indirectly through the RR interval series, characterized by its trajectory, the stability structure of its deterministic drift, persistent state regions, and transitions. This builds on the neurovisceral integration framework (Thayer and Lane 2000) and recognizes that cardiac autonomic control is coupled to central brain states, metabolic demands, inflammatory pathways, behavior, and the individual’s exposome and clinical status (Wild 2005). Complementary approaches exist: nonlinear HRV analyses quantify temporal complexity; state-space models estimate latent cardiovascular dynamics (Rosas et al. 2024; Castillo-Aguilar et al. 2026); network physiology examines interactions among organ systems; dynamical resilience frameworks characterize recovery and critical transitions; and landscape and escape-rate formalisms from nonequilibrium statistical physics supply the mathematics of multistable stochastic systems (Ao 2004; Wang, Xu, and Wang 2008; Hänggi, Talkner, and Borkovec 1990). What is missing is a single structure that separates latent autonomic state, deterministic landscape, stochastic trajectory, perturbation response, and dynamic phenotype, and that states what can and cannot be inferred about each.

In this article we develop such a structure, the Systems Theory of Cardiac-Autonomic Trajectories and Equilibria (STATE). It is a theoretical and computational study, not an empirical validation. It offers a minimal set of postulates, definitions, assumptions, and non-assumptions from which specific models can be derived and tested; two formal results that bound what the theory can claim (Section 4 and Appendix A); and a deliberately small simulation (Section 6) showing that recovery descriptors carry information that baseline and peak summaries do not. The framework does not assume that latent trajectories are uniquely recoverable from every RR recording; identifiability and inferential adequacy are treated as properties to be established for each implementation. Section 2 presents the framework and its postulates, Section 3 its formal core, Section 4 and Section 5 its inferential limits, Section 6 the illustration, and Section 7 and Section 8 the testable hypotheses, limitations, and conditions under which we would abandon the theory.

## 2 The STATE framework

STATE proposes that cardiac autonomic regulation is a latent dynamical process, bidirectionally coupled (rather than linearly caused) to a broader system of physiological, behavioral, molecular, and environmental domains. The autonomic component is represented as a low-dimensional latent state, *A*_*t*_ ∈ ℝ^*d*^, observed only indirectly through the RR interval series: a representation of the information in autonomic regulation relevant to cardiac dynamics, not a claim about the number or identity of the physiological variables that generate it. This reflects the theory’s central epistemological commitment: a physiological system is not its latent representation, and a latent representation is not what can be observed from it or inferred about it,

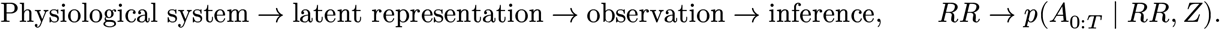

STATE is accordingly not a theory that equates measured cardiac signals with autonomic activity. Three distinctions follow and recur throughout: a latent state is not the complete physiological state; a model parameter is not, by itself, a biological mechanism; and an identified mathematical structure is not a causal structure.

We adopt a two-dimensional instantiation,

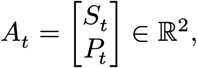

an interpretable decomposition into two coordinates associated with sympathetic and parasym-pathetic cardiac modulation. These are a low-dimensional representation of autonomic influences on cardiac dynamics, not direct measurements of neural activity, and the two-dimensional choice is a modeling hypothesis, not a claim that the physiological system has two degrees of freedom.

The trajectory *A*(*t*) traces how this state evolves, its structure set jointly by the deterministic drift and stochastic fluctuations. We call the resulting context-dependent collection of state regions, stability properties, transitions, and perturbation responses the *autonomic dynamical landscape*: a compact description of the drift’s vector field together with the fluctuations and transitions around it, not a literal surface or a scalar potential *V* (*A*), which the theory neither requires nor assumes. The landscape does not exist independently of the organism: the broader system’s state can shape autonomic dynamics and, reciprocally, autonomic dynamics can be associated with changes elsewhere, in a chain of couplings

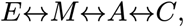

where *E* is the exposome, *M* the multidomain state, *A* the cardiac autonomic state, and *C* the clinical or functional state, couplings that are context-, time-, and state-dependent rather than uniformly causal, in the spirit of the allostatic-load (McEwen and Stellar 1993) and network-physiology and network-medicine (Bartsch et al. 2015; Barabási, Gulbahce, and Loscalzo 2011) literatures (Figure 1).

**Figure 1:**
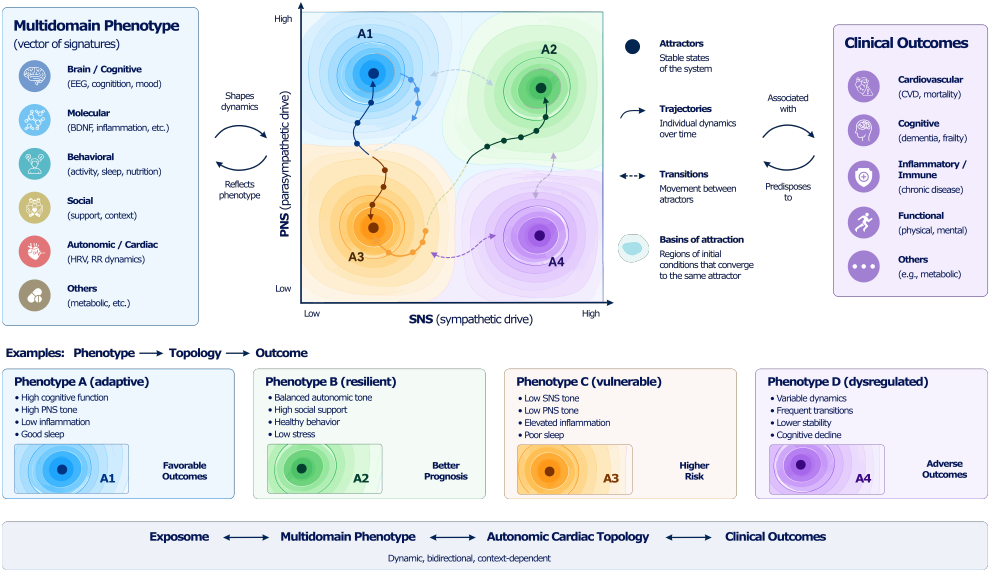
Schematic representation of STATE, coupling the dynamic exposome (*E*), the multidomain state (*M*), the cardiac autonomic state (*A*), and the clinical state (*C*). The attractors A1–A4, their basins, and the phenotype–outcome pairings shown are hypothetical illustrations of configurations the theory admits; STATE does not assume that any attractor is favorable or unfavorable, nor that the depicted associations are causal (Section 5.2).

### 2.1 Foundational postulates

STATE rests on four postulates, which state what the theory affirms about the structure of cardiac autonomic regulation, as distinct from the auxiliary conditions needed to test it, which are stated in Section 5.1. Postulates 1, 3, and 4 are empirical claims, not established properties of autonomic physiology; Postulate 2 is a methodological principle, justified only to the extent that the dynamical descriptors it calls for carry information that static summaries do not (hypotheses *H*_1_ and *H*_4_).

#### Postulate 1 (Low-dimensional representation)

A relevant component of cardiac autonomic regulation can be represented by a low-dimensional latent state whose temporal evolution contains information that is not recoverable from isolated summary statistics alone.

#### Postulate 2 (Dynamics over location)

The characterization of an autonomic state should depend not only on its location in latent state space but also on the dynamical properties governing its persistence, transition, and response to perturbation.

#### Postulate 3 (Context-dependence)

The dynamical landscape is context-dependent: changes in multidomain, environmental, or clinical conditions may alter the drift and stochastic structure governing autonomic trajectories.

#### Postulate 4 (Individual variation)

Individuals may differ not only in their mean autonomic state but also in the geometry and stochastic properties of their dynamical landscapes.

Together, these postulates imply that an individual’s physiological phenotype cannot be reduced to a static set of values: two individuals may share the same mean RR interval, RMSSD, or estimated autonomic state and yet differ in the stability structure of their dynamics, the persistence of state regions, their transition probabilities, and their recovery from a challenge. We hold that these properties may carry physiological information that is lost in static summaries, and we treat this as a claim to be tested, not a premise. Postulate 3 would additionally require within-person comparisons across contexts, which *H*_1_ to *H*_4_ do not cover and which we leave to future work.

### 2.2 Resilience as a dynamical property

We understand resilience as the response of a system’s trajectory to perturbation, not as a fixed trait, echoing the use of the term in dynamical-systems research since its introduction in ecology (Holling 1973). A rapid return to baseline is not necessarily synonymous with greater resilience; adaptive reconfiguration to a new state may also be an appropriate response. Resilience is studied through challenge protocols such as graded exercise, head-up tilt, the cold pressor test, or the Valsalva maneuver. The view is consistent with critical slowing down as an early-warning signal in complex systems, including human physiological and mood time series (Scheffer et al. 2009; Leemput et al. 2014), with dynamical resilience indicators in aging and geriatric populations (Gijzel et al. 2017; Rector et al. 2021), and with a logic long established in cardiology, where the rate of heart-rate decline after exercise is itself a validated prognostic marker of recovery capacity (Cole et al. 1999; Qiu et al. 2017). Formal descriptors are defined in Section 3.3.

## 3 Formal core

### 3.1 Stochastic formulation and observation model

Multidomain, exposomic, and clinical influences enter the core autonomic subsystem as a context variable, *η*_*t*_ = (*M*_*t*_, *E*_*t*_, *C*_*t*_), that shapes autonomic dynamics without itself requiring a fully estimated dynamical model. The autonomic state then evolves as a stochastic differential equation (SDE),

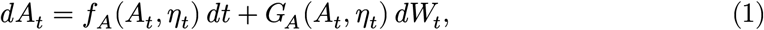

which separates a deterministic component, *f*_*A*_(*⋅*) *dt*, describing the expected local direction of change, from a stochastic component, *G*_*A*_(*⋅*) *dW*_*t*_, describing process variability not captured by the drift. This variability is conceptually distinct from measurement error: it may reflect genuine physiological fluctuation, unobserved inputs, or model approximation. In extended formulations, components of *η*_*t*_ may themselves be modeled as dynamic states coupled to *A*_*t*_, but this is warranted only when the temporal resolution, measurement structure, and scientific question justify it.

For the cardiac signal, the generic observation model is

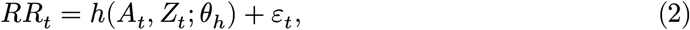

where *Z*_*t*_ denotes additional observed physiological variables relevant to cardiac timing (such as respiration), *θ*_*h*_ the parameters of the observation mapping, and *ε*_*t*_ measurement or residual error, distinct from the process variability above. The inferential target is the posterior *p*(*A*_0:*T*_, *θ* ∣ *RR*_0:*T*_, *Z*_0:*T*_) or its marginal over the latent trajectory. A simple admissible instance is the linear, beat-synchronous observation 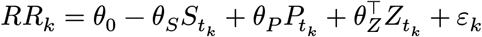 with *t*_*k*+1_ = *t*_*k*_ + *RR*_*k*_, so that the sampling times depend on the latent state; here *θ*_*S*_, *θ*_*P*_ > 0 under the convention that larger *S*_*t*_ (*P*_*t*_) shortens (lengthens) the RR interval. The theory does not require this instance, and the illustration of Section 6 deliberately works with the latent state itself, not with an RR series.

The latent trajectory is therefore a probabilistic object and not a measured signal. Disentangling biological stochasticity from observation error is a non-trivial inverse problem, and any inferred landscape depends on the adequacy and identifiability of the chosen models, so derived quantities should be interpreted together with their inferential uncertainty.

### 3.2 Geometry of the autonomic state space

For a fixed or slowly varying context *η*, the drift defines a deterministic skeleton, *dA*/*dt* = *f*_*A*_(*A, η*), from which equilibria *A*^*^(*η*) (with *f*_*A*_(*A*^*^, *η*) = 0), basins of attraction 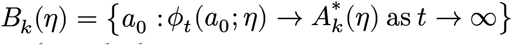 (with *ϕ*_*t*_ the deterministic flow), and local stability through the eigenvalues of the Jacobian 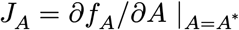 can be defined, in the tradition that links bifurcations in the deterministic part of physiological control models to qualitative changes in dynamics associated with disease (Mackey and Glass 1977). We stress that *f*_*A*_ is a model component, not a hypothesized physiological mechanism: it may represent the aggregate resultant of many unobserved regulatory processes.

The stochastic system must be interpreted separately. Because of the diffusion term, an observed trajectory need not converge to a deterministic equilibrium: it may fluctuate around stable regions, leave them, or move between them. We therefore distinguish deterministic attractors of the drift from stochastic state regions characterized by persistence, occupancy, and transition probabilities, such as the finite-horizon transition probability *P*_*jk*_(*H*) = Pr(*A*_*t*+*H*_ ∈ *S*_*k*_ ∣ *A*_*t*_ ∈ *S*_*j*_, *η*) between regions *S*_*j*_ and *S*_*k*_. An attractor is a property of the deterministic skeleton, whereas a persistent state region is a property of the realized process; the two need not coincide, and STATE does not require them to.

### 3.3 Resilience descriptors

Resilience is formalized here as the response of the autonomic trajectory to an explicit perturbation, building on the state regions and transition probabilities introduced above.

Let *ℛ*_*rest*_ denote a baseline resting state region in autonomic space rather than a single fixed point. Following an applied physiological perturbation *u*(*t*) (e.g., the onset of ergometer exercise or orthostatic stress; the symbol *u* is used to avoid confusion with the parasympathetic coordinate *P*_*t*_), the system moves away from this baseline region, enters a perturbed regulatory regime, and subsequently returns toward the baseline region or transitions to an altered persistent state during recovery:

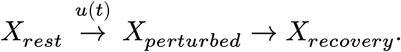

More generally, the perturbation can be incorporated directly into the core autonomic dynamics as an explicit forcing term, *dA*_*t*_ = *f*_*A*_(*A*_*t*_, *η*_*t*_, *u*_*t*_) *dt* + *G*_*A*_(*A*_*t*_, *η*_*t*_) *dW*_*t*_.

For a suitable distance function *d*(*⋅, ℛ*_*rest*_) defined on a specified and reproducible state-space metric, the maximum autonomic displacement quantifies the magnitude of systemic departure from baseline,

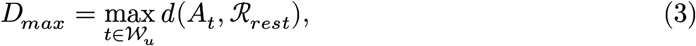

where *W*_*u*_ denotes the active perturbation window, whose end is denoted *t*_*end*_. Displacement is meaningful relative to the chosen parameterization and metric of the latent state space.

Membership in the resting region is assessed with a stability tolerance *δ*: we write 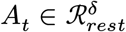 when *d*(*A*_*t*_, *ℛ*_*rest*_) ≤ *δ*. Recovery time is measured from the end of the perturbation window and is the earliest delay after which the trajectory remains within the tolerance region for a dwell time *τ*,

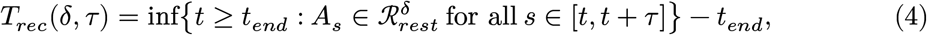

with *T*_*rec*_ = ∞ (right-censored at the observation horizon) when no such *t* exists. For a specified observation horizon *H*, the probability of sustained return to the resting regime and the probability of a regime shift are

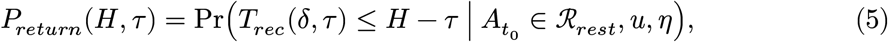

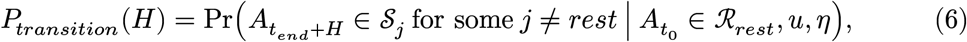

where *S*_*j*_ are persistent state regions other than *ℛ*_*rest*_ (the transition probabilities introduced above, applied to the post-perturbation state). Taken together, these quantities characterize resilience as a multidimensional response profile rather than a single score,

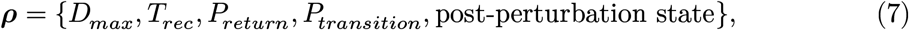

where the post-perturbation state records which region, if any, the system occupies at the end of the horizon. Each component reflects a distinct aspect of neurocardiac control. Under this formulation, “autonomic resilience” describes the stochastic temporal behavior of the neurocardiac system in response to physiological challenge, rather than the baseline magnitude of vagal tone or HRV.

Three specification rules follow from the stochastic character of the process. First, *δ, τ*, and *H* are part of the definition and must be reported with every descriptor: for a positive Harris recurrent diffusion (for instance, one with coercive drift and nondegenerate noise), every open region is entered with probability one as *H* → ∞, so *P*_*return*_(*H, τ*) → 1 for any *τ*, and “failure to return” is always a statement about a finite horizon. Second, *δ* should be calibrated from the resting regime alone, independently of the perturbation response and of the outcome (the third assumption in Section 5.1); a convenient rule is a high quantile (here 0.999) of *d*(*A*_*t*_, *ℛ*_*rest*_) in a resting segment of the same recording. Third, when *ℛ*_*rest*_ is identified with a basin of the deterministic skeleton, *δ* must keep 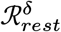 inside that basin, and *T*_*rec*_ can detect a transition to another regime only if *δ* is smaller than the distance between that regime and *ℛ*_*rest*_.

### 3.4 Dynamic phenotype and clinical outcomes

Beyond the autonomic state itself, the theory carries a multidomain profile *M*_*t*_ (brain, cognitive, molecular, immune, behavioral, social, and further components) with its own dynamics, so that the relationship *A*_*t*_ ↔ *M*_*t*_ is bidirectional and not a single causal direction. Combining the autonomic trajectory with this profile, a compact representation of individual *i*’s dynamic phenotype is

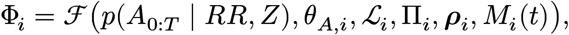

where *p*(*A*_0:*T*_ ∣ *RR, Z*) is the posterior over the latent trajectory, *θ*_*A*,*i*_ the parameters governing the individual’s dynamics, ℒ_*i*_ the context-dependent landscape, Π_*i*_ the transition structure between persistent regions, *ρ*_*i*_ the resilience profile, and *M*_*i*_(*t*) the multidomain profile. The dynamic phenotype is thus not a single latent score but a probabilistic representation of how an individual’s autonomic system evolves, persists, transitions, and responds to perturbation. Clinical outcomes are modeled as *O*_*i*_ *∼ p*(*O*_*i*_ ∣ Φ_*i*_, *E*_*i*_, *C*_*i*_): they depend probabilistically on the phenotype and on other factors, and the theory does not require that all outcome variation be explained by the autonomic system.

## 4 What can be inferred from RR intervals

Because STATE represents autonomic regulation through latent variables, the interpretation of the resulting landscape depends on the identifiability of the chosen representation from the available observations. Recent generative state-space approaches already make this concern explicit by reporting full posterior uncertainty over latent heart-rate trajectories (Rosas et al. 2024; Castillo-Aguilar et al. 2026). The theory does not assume that a single RR series can always be decomposed uniquely into sympathetic- and parasympathetic-associated coordinates: identifiability is a property of the observation model, the stochastic dynamics, the available covariates, the recording duration, and the experimental design, and spans structural and practical identifiability (Wieland et al. 2021).

### Proposition 1

(identifiability of the sympathetic and parasympathetic coordinates from a scalar output)

Let *dA*_*t*_ = *F A*_*t*_ *dt* + *σ dW*_*t*_ with *F* ∈ ℝ^2×2^ Hurwitz and *W* a two-dimensional standard Brownian motion, observed through *y*_*t*_ = *c*^⊤^*A*_*t*_ + *ε*_*t*_ with *c* ≠ 0 and *ε*_*t*_ measurement error independent of *W* .

i. If *F* = −*λI* (isotropic, uncoupled dynamics), then for every *c* the law of *y* depends on the dynamics only through (*λ, σ*‖*c*‖). The component of *A*_*t*_ orthogonal to *c* is an Ornstein– Uhlenbeck process independent of *y*, so its posterior equals its prior and the coordinates (*S*_*t*_, *P*_*t*_) are not identifiable, however long the recording.
ii. If *F* = diag(−*λ*_*S*_, −*λ*_*P*_) (uncoupled dynamics with distinct rates), the pair (*F*, *c*^⊤^) is observable if and only if *c*_*S*_*c*_*P*_ (*λ*_*S*_ − *λ*_*P*_) ≠ 0.
iii. If 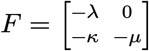 (the structure of System B linearized at rest in Section 6), the pair (*F*, *c*^⊤^) is observable if and only if *c*_*P*_ [*c*_*S*_(*λ* − *µ*) + *c*_*P*_ *κ*] ≠ 0.

The proof, a symmetry argument for (i) and the Kalman rank condition for (ii) and (iii), is given in Appendix A.

Observability in the sense of (ii) and (iii) guarantees that the latent state can in principle be reconstructed, not that it is reconstructed with adequate precision: both determinants vanish continuously as the rates approach each other (and, in (iii), as the coupling vanishes), so near-isotropic dynamics are practically unidentifiable even when formally observable, and practical identifiability always depends on the signal-to-noise ratio and the recording duration. Proposition 1 therefore makes precise why the two-dimensional instantiation of STATE requires, to be interpretable, distinct time scales, coupling between the coordinates, additional observed signals *Z*_*t*_ such as respiration, or perturbation protocols; without them, *S*_*t*_ and *P*_*t*_ would be labels without identifiable content. The two autonomic limbs are commonly described as acting on different time scales, which, if so, favors case (ii), but this must be established for each implementation. The systems of Section 6 are coupled; for *c* = (−1, 1) and (*λ, µ, κ*) = (0.274, 0.48, 0.24) the determinant in (iii) is 0.446 ≠ 0, whereas their uncoupled isotropic counterpart (*κ* = 0, *λ* = *µ*) would have rank one for every *c*.

Accordingly, a STATE implementation should distinguish three levels of inference. The first is reconstruction of the latent trajectory, expressed through *p*(*A*_0:*T*_ ∣ *RR, Z*). The second is estimation of dynamical properties such as local stability, state persistence, or transition probabilities. The third is interpretation of these properties in physiological or clinical terms. Uncertainty at the first level propagates to the subsequent levels and should not be ignored when defining individual dynamic phenotypes.

This establishes an explicit inferential constraint on the theory: a dynamical feature should not be interpreted as an individual physiological property unless it is sufficiently identifiable and robust to reasonable changes in model specification, observation assumptions, and measurement error. The purpose of STATE is therefore not to guarantee recovery of a unique latent autonomic state from every recording, but to provide a formal framework in which the adequacy of such reconstruction can be empirically evaluated.

## 5 Assumptions and boundaries of the theory

### 5.1 Assumptions

The postulates state what STATE affirms; testing them against data requires a further, narrowly methodological set of assumptions. First, the RR series is assumed to carry information sufficient to constrain at least some dimensions of the latent state, without implying that the mapping from *A*_*t*_ to *RR*_*t*_ is invertible. Second, the observation model *h*(*⋅*) and the drift and diffusion functions are assumed to be adequate approximations over the recording window and protocol; misspecification can bias the inferred trajectory and every descriptor computed from it. Third, the state regions, the tolerance *δ*, and the dwell time *τ* must be specified independently of the outcome under study, to avoid circularity between the definition of recovery and its association with an endpoint (a resting-segment calibration rule is given in Section 3.3). Fourth, persistence and transition descriptors are estimable only if the recording is long relative to the relaxation times of the landscape and, in multistable landscapes, comparable to the escape times between regions or accompanied by an explicit perturbation protocol; otherwise baseline summaries describe a transient and not the stationary behavior of the process (Appendix A). Finally, comparisons across individuals or conditions assume sufficiently standardized perturbation protocols, durations, and covariates. None of these is guaranteed in a given dataset; each is a condition for the postulates to be tested validly and can in principle be checked empirically.

### 5.2 Boundaries

The boundaries of the theory matter as much as its content. STATE is not a direct measurement model: it does not assume that RR measures SNS or PNS activity, that the latent state is uniquely identifiable from a single recording, or that inferential uncertainty can be ignored. It is not a fixed autonomic-balance model: it requires neither a difference (*SNS* − *P NS*) nor a ratio (*SNS*/*P NS*) as the system’s definition, since its object is the joint dynamic space. It is not a deterministic attractor theory: attractors and basins belong to the deterministic skeleton, whereas the stochastic process is characterized by persistence, fluctuation, and transition. It does not assume a “healthy attractor” (a structure can be stable and still maladaptive) or a common landscape for all individuals. And it is not a causal theory: observing *A* ↔ *M* does not license *A* → *M*, and establishing causality requires independent experimental designs or formal causal inference. A latent coordinate, state region, or transition pattern is an inferential construct whose physiological interpretation requires convergent evidence from independent measurements or manipulation. The guiding principle is *Brain* ↔ *Heart* within a broader physiological network, and the autonomic system is one determinant among several of any clinical outcome. Finally, the theory does not equate resilience with rapid return to baseline, does not require all domains to be modeled as latent processes at once, and does not require continuous-time state-space modeling as its only implementation.

## 6 A minimal illustration: similar baseline and peak, different fate

To show why we hold that recovery descriptors carry information that baseline and peak summaries do not, we use the smallest model that can make the point: two systems whose baseline variability and peak response to a perturbation are comparable but whose dynamical organization differs. They are built to differ only in the nonlinearity of the sympathetic drift, so that differences in outcome cannot be attributed to stiffness at rest, coupling, or noise, but only to a second attractor. The bistable system echoes evidence that the deterministic component fitted to real human heart-rate fluctuations can itself be bistable (Kuusela, Shepherd, and Hietarinta 2003). Both systems are specified by hand and no parameter was estimated from data; the full specification is in Appendix B.

System A is monostable and is exactly the linearization of System B at rest: both have the same Jacobian at the origin, the same coupling, and the same noise. System B has a stable resting attractor, a saddle, and a stable stress attractor, with *ℛ*_*rest*_ inside the resting basin. The sympathetic coordinate of System B is an autonomous one-dimensional diffusion in a symmetric double-well potential, for which exact results are available (Appendix A): its stationary law places half of its mass in each basin and has 17 times the variance of System A, the mean escape time from rest is 217 s at the nominal noise (against a 25-s baseline), and a constant forcing displaces it 5.2 times more easily than it displaces System A to the same location. An identical transient pulse, 15.8 times the critical forcing of System B, was applied to both systems in 2000 independent replicates each, and the descriptors of Section 3.3 were computed with a tolerance *δ* = 150.7 ms fixed a priori from the resting regime of System A.

During the 25-s baseline both systems fluctuate around *ℛ*_*rest*_ with comparable variability (median SD of *d* of 22.3 ms in System A and 23.0 ms in System B; Table 2). The area under the curve (AUC) for discriminating B from A on this descriptor was 0.56 (95% bootstrap interval 0.54–0.57), against 0.5 for no separation. The distribution of System B has a heavier upper tail (97.5th percentile 85 ms against 37 ms) because in 7.6% of its replicates the sympathetic coordinate crossed the saddle during the baseline. Under the pulse, both systems are displaced to nearly the same peak (median *D*_*max*_ of 725 ms in A and 717 ms in B; AUC 0.42, 0.40–0.44): peak displacement did not discriminate between them at this pulse amplitude (see Robustness). The fate after the pulse did. System A returned to 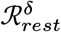 and remained there for at least 5 s in all replicates (median *T*_*rec*_ 6.2 s, 2.5th–97.5th percentile 4.4–9.6 s), whereas System B did so in only 6.7% of replicates (95% interval 5.6–7.8%), so that *T*_*rec*_ was right-censored at the 55-s horizon in 93.3% of them, and 91.2% (89.9–92.4%) of the replicates occupied the basin of the stress attractor at the end of the horizon (Figure 2). The recovery descriptors separated the systems almost perfectly (AUC for *T*_*rec*_ of 1.00, which largely reflects the right-censoring of System B, so the proportions above are the more informative summaries), whereas baseline variability and peak displacement did not at this pulse amplitude.

**Figure 2:**
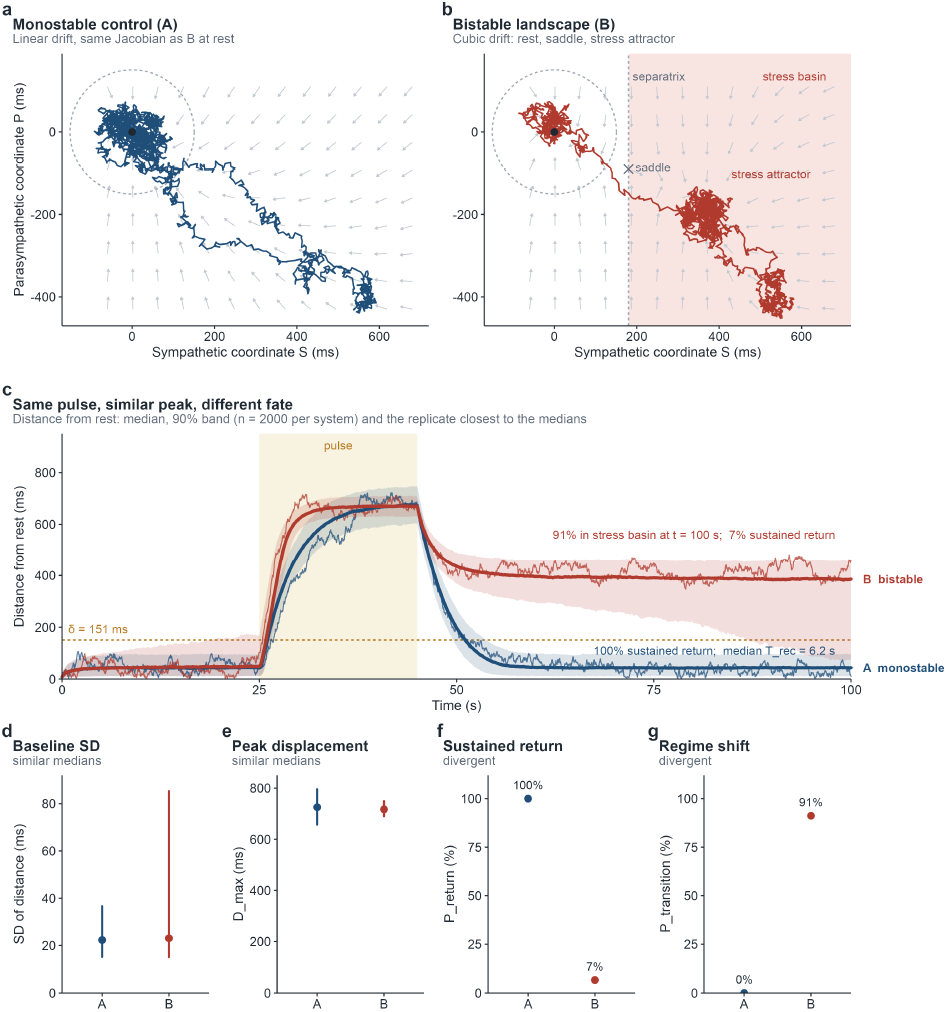
Controlled contrast between a monostable (A) and a bistable (B) autonomic landscape under an identical pulse. (a, b) State-space trajectories of the replicate closest to the ensemble medians, with the vector field of the deterministic drift and the resting tolerance region (gray disc of radius *δ*); in (b), the separatrix, saddle (×), stress attractor (▴), and the basin of the stress attractor (shaded). (c) State distance from *ℛ*_*rest*_: median and 90% band over 2000 replicates and the same representative replicates; the dashed line is *δ*. (d, e) Baseline SD and *D*_*max*_: median with 2.5th–97.5th percentile interval. (f, g) Proportion of replicates with sustained return *P*_*return*_(*H, τ*) and in the stress basin at *t* = 100 s, *P*_*transition*_(*H*), with 95% Wilson intervals.

Three conditions delimit the result, and they are the lessons we draw from it (robustness analyses in Appendix B). First, baseline equivalence holds only while the baseline window is short relative to the escape time and the noise is small relative to the barrier; at higher noise the two systems become distinguishable at baseline and the contrast degrades as the exact results predict. Second, the coincidence of peaks is specific to the pulse amplitude used: across amplitudes the bistable system’s peak displacement is up to 2.8 times larger at intermediate amplitudes, and the two coincide only at the largest one, so peak displacement cannot by itself identify the regime. Third, recovery descriptors inherit the analyst’s specification: the tolerance must lie inside the resting basin yet below the distance of the alternative regime, the horizon is finite by necessity, and the noise regime determines whether “failure to return” is a property of the system or of the realization.

The illustration does not show that human autonomic dynamics are bistable, that these descriptors can be recovered from RR recordings (the latent distance simulated here is not an RR series, and its standard deviation is not SDNN), or what effect sizes to expect in real data. The time scales (relaxation times of 2–4 s for both coordinates) are illustrative and were not calibrated to autonomic physiology, in which the vagal limb is generally considered faster than the sympathetic; the recovery times should not be read as physiological estimates. What it does show, in a system whose ground truth is known, is that descriptors computed at baseline, and peak displacement at a given amplitude, need not distinguish two dynamical organizations, whereas the fate after a perturbation can.

## 7 Testable hypotheses and potential applications

The framework suggests applications in four areas, each an empirical hypothesis and not a pre-supposition. In physiological phenotyping, individuals would be described by latent trajectories, stability, persistence, transitions, and perturbation response, not solely by point measures such as

HR, RMSSD, or SDNN; the open question is whether such descriptors add information. In stress and resilience testing, a standardized perturbation such as exercise becomes a perturbation test of the system, interpreted through the landscape and not through peak output alone, pending validation against real recordings. In aging and disease, one can ask whether changes in the number, stability, or transition structure of state regions, or in perturbation response, accompany aging or particular clinical outcomes, consistent with dynamical resilience indicators in geriatric research (Gijzel et al. 2017; Rector et al. 2021), with declining physiologic complexity in aging and disease (Costa, Goldberger, and Peng 2002; Jandackova et al. 2016), with reduced HRV in depression (Kemp et al. 2010), and with slowed physiological recovery as a marker of impending deterioration (Olde Rikkert et al. 2016), without assuming in advance that any configuration is pathological. In personalized modeling, an individual landscape ℒ_*i*_ can be estimated for each person and related to clinical evolution, in parallel with digital phenotyping (Onnela and Rauch 2016) and N-of-1 designs (Percha et al. 2019); continuous HRV monitoring in free-living conditions makes such longitudinal designs increasingly feasible (Gaur et al. 2024). Because the context variable *η*_*t*_ may include respiration, blood pressure, inflammatory markers, EEG, or behavioral context, multimodal integration does not require collapsing these into a composite score, although jointly estimating them increases dimensionality and data requirements.

The framework generates four directly testable hypotheses that define the empirical program needed to evaluate STATE:

*H*_1_ : dynamic descriptors contain information not captured by static HRV summaries;

*H*_2_ : similar baseline HRV does not imply similar perturbation-response dynamics;

*H*_3_ : dynamic phenotype descriptors are reproducible within individuals across repeated assessments;

*H*_4_ : specific dynamical properties predict subsequent physiological or clinical states.

Each can be evaluated by applying state-space inference implementations (Rosas et al. 2024; Castillo-Aguilar et al. 2026) to recordings acquired under standardized perturbation protocols and repeated or longitudinal assessments. Table 1 states, for each hypothesis, the estimand, the minimal design, and the result that would count against it. In every case the specification (*δ, τ*, *H*, the distance metric, and the perturbation protocol) and the minimal useful difference are registered before analysis, and the descriptors are carried with the uncertainty of the latent reconstruction (Section 4). A rest–exercise–rest protocol is a natural design: it provides a resting segment from which *δ* can be calibrated, a perturbation of known timing, and a recovery segment in which *T*_*rec*_, *P*_*return*_, and *P*_*transition*_ are observed.

**Table 1:** Operational form of the four hypotheses.

| Hypothesis | Estimand | Minimal design | Not supported if |
|---|---|---|---|
| *H*<sub>1</sub> | Incremental information: difference in cross-validated performance (e.g., AUC or log-likelihood) for a physiological or clinical endpoint between a model with static HRV summaries plus **ρ̂** and the same model without **ρ̂** | Cohort with a standardized perturbation and a follow-up endpoint; nested models | The 95% interval of the difference contains no value above the minimal useful difference |
| *H*<sub>2</sub> | Discordance at matched baseline: dispersion of **ρ̂** among pairs matched on static HRV within a tolerance *ϵ*, relative to its dispersion across repeated assessments of the same individual | As *H*<sub>1</sub>, with at least two assessments per individual | Between-individual dispersion in matched pairs is not larger than within-individual dispersion |
| *H*<sub>3</sub> | Reproducibility: intraclass correlation (or concordance) of each descriptor across repeated assessments under the same protocol | At least two assessments separated by a period over which no change is expected | The lower bound of the 95% interval of the correlation is below the pre-specified threshold |
| *H*<sub>4</sub> | Prediction of subsequent states: incremental discrimination and calibration for a future physiological or clinical state given **ρ̂**, beyond static HRV and clinical covariates | Longitudinal cohort with a baseline protocol and a follow-up | Out-of-sample discrimination and calibration do not improve beyond the minimal useful difference |

**Table 2:** Descriptors in 2000 replicates per system (*σ* = 0.10, *δ* = 150.7 ms, *τ* = 5 s, *H* = 55 s). Continuous descriptors: median [2.5th, 97.5th percentile]; percentages: proportion [95% Wilson interval]. 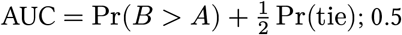 indicates no separation.

| Descriptor | System A<br>(monostable) | System B<br>(bistable) | AUC, B vs A<br>(95% CI) |
|---|---|---|---|
| Baseline SD of $d(A_t, \mathcal{R}_{rest})$, $t < 25$ s (ms) | 22.3 [15.2, 36.6] | 23.0 [15.1, 85.3] | 0.56 (0.54–0.57) |
| Baseline windows with a smoothed excursion beyond $\delta$ (%) | 2.8 [2.2, 3.6] | 14.7 [13.2, 16.3] |  |
| Saddle crossed during baseline (%) | 0.0 [0.0, 0.2] | 7.6 [6.5, 8.8] |  |
| Peak displacement $D_{max}$ (ms) | 725.2 [656.8, 796.1] | 717.1 [690.3, 749.1] | 0.42 (0.40–0.44) |
| Sustained return $P_{return}(55\,\text{s}, 5\,\text{s})$ (%) | 100.0 [99.8, 100.0] | 6.7 [5.6, 7.8] |  |
| Recovery time $T_{rec}(\delta, 5\,\text{s})$, median (s) | 6.2 | 55.0 (right-censored) | 1.00 |
| Regime shift $P_{transition}(55\,\text{s})$ (%) | 0.0 [0.0, 0.2] | 91.2 [89.9, 92.4] |  |

## 8 Discussion

We offer STATE as the foundation of a refinable theory, not a finished model. Its core is a small number of postulates (Section 2.1), a formal vocabulary that keeps the physiological system, its latent representation, its observation, and the inference drawn from it distinct (Section 3), and an explicit statement of what the theory does and does not assume (Section 5). What it contributes is a way of formulating questions about autonomic regulation so that they can be answered, or refuted, with data: which properties belong to the deterministic drift and which to the realized stochastic process, which quantities can be identified from a given recording, and which descriptors of perturbation response are expected to carry information beyond static HRV summaries.

STATE is intended to complement existing frameworks. Neurovisceral integration (Thayer and Lane 2000), allostasis (McEwen and Stellar 1993), and network physiology (Bartsch et al. 2015) describe how regulation is distributed across brain, autonomic, and peripheral systems, but do not prescribe a state-space description of the cardiac autonomic component or a way of characterizing its stability and perturbation response. Research on critical transitions and dynamical resilience (Scheffer et al. 2009; Leemput et al. 2014; Gijzel et al. 2017) shows that recovery carries information about proximity to a change of state; STATE situates such indicators within an explicit driftdefined landscape, so that slowing recovery, displacement, or a transition between regions can be read against a stated model and not as isolated indices. Landscape and flux formulations (Ao 2004; Wang, Xu, and Wang 2008) and escape-rate theory (Hänggi, Talkner, and Borkovec 1990) supply the mathematics of multistable stochastic systems; what STATE adds is coupling to an observation-limited, perturbation-based physiological protocol and an account of which quantities are identifiable. Because autonomic drift fields need not be gradient fields (System B of Section 6 is not), STATE speaks of basins of the deterministic skeleton and persistent stochastic regions and not of a scalar potential. Generative state-space approaches (Rosas et al. 2024; Castillo-Aguilar et al. 2026) are candidate implementations of the inferential layer, and earlier fits that revealed bistable deterministic components in human heart-rate fluctuations (Kuusela, Shepherd, and Hietarinta 2003) are consistent with the multistable landscapes STATE admits, although they do not show that this organization is general.

A fourth lesson of the illustration concerns protocol design: a bistable landscape can be dislodged by perturbations several times smaller than those that displace its linear counterpart to the same location (by a factor of 3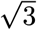 for the symmetric cubic), so a protocol of fixed amplitude may saturate the response and reveal nothing about graded vulnerability. Graded protocols are there-fore preferable, and the proportion of replicates shifting regime differed between the systems (0% against 22–92%) at every amplitude tested, which supports describing resilience through recovery and regime shift and not through the amplitude of the response alone.

Several limitations bound the present formulation. First, the theory has not been evaluated on empirical recordings: the illustration shows that descriptors of perturbation response can separate landscapes with comparable baseline variability and peak response when ground truth is known, but not that human autonomic dynamics are organized in this way. Second, the two-dimensional representation is a modeling hypothesis, and the mapping from the latent coordinates to sympathetic and parasympathetic physiology is interpretive rather than identified. Third, identifiability from RR intervals alone may be limited: for isotropic uncoupled dynamics the two coordinates cannot be separated from a scalar signal, so coupling structure, additional signals such as respiration, or perturbation protocols are required (Section 4). Fourth, the resilience descriptors depend on analyst-specified choices (the distance metric, *δ, τ*, *H*, and the protocol) and are comparable only under shared specifications (Section 3.3). Fifth, the exact results are for minimal systems; their extension to state-dependent diffusion, higher dimensions, and non-gradient multistable fields, where escape rates depend on a quasi-potential and a curl flux (Ao 2004; Wang, Xu, and Wang 2008), is not attempted. Finally, extending the state to multidomain dynamics increases dimensionality and data requirements, and the theory does not prescribe when that is warranted.

Because postulates and boundaries are explicit, the theory can be refined along identifiable axes and not revised as a whole: the latent dimension can grow when it demonstrably improves inferential adequacy or prediction; the drift and diffusion can be specialized from linear-Gaussian processes to nonlinear drifts with multiple equilibria and state-dependent diffusion; the context variable can become a dynamic component; and the definitions of regions, transitions, and resilience descriptors can be tightened or replaced as empirical work shows which are reproducible and informative. Each refinement leaves intact the separation between the physiological system, its representation, its observation, and the inference drawn from it.

The theory is also open to refutation. If dynamic descriptors add no information beyond static HRV summaries (*H*_1_), if matched baseline HRV implies matched perturbation-response dynamics (*H*_2_), if the descriptors are not reproducible within individuals (*H*_3_), or if they do not predict subsequent physiological or clinical states (*H*_4_), then the added complexity of a latent dynamical description would not be justified for the corresponding application, and the postulates concerning the informational value of dynamics would need to be revised or restricted. Conversely, support for these hypotheses would motivate the more demanding program of validating specific implementations under realistic recording conditions.

## 9 Conclusions

The central proposition of STATE is that cardiac autonomic regulation can be represented as a latent stochastic dynamical system embedded within a broader multidomain physiological context:

Exposome ↔ Multidomain physiology ↔ Cardiac autonomic dynamics ↔ Clinical state .

Within the autonomic component, the inferential chain is

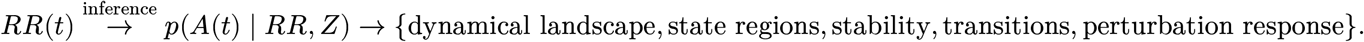

The resulting dynamic phenotype represents the individual’s latent autonomic organization together with the uncertainty associated with its reconstruction and the contextual factors that shape its dynamics.

The central conceptual distinction of the theory is between the deterministic drift and the stochastic trajectory. The drift defines the context-dependent dynamical landscape in which equilibria, local stability, and deterministic basins can be studied, whereas the stochastic component determines fluctuations, persistence, escape, and transitions between regions of the landscape. This formulation permits the study of resilience without requiring recovery to a single fixed point. It also locates the fundamental object of STATE not in HRV itself, nor in a single measure of autonomic balance, but in the inferred dynamical organization of cardiac autonomic regulation and its interaction with the broader physiological system. RR intervals provide an accessible observation through which this latent organization may be estimated, while state-space and stochastic modeling provide the computational tools for doing so. The computational illustration in Section 6 shows the practical consequence: two landscapes with the same linearization at rest, comparable baseline variability, and, at the amplitude used, comparable peak displacement can differ in their fate after a common perturbation, a difference that only descriptors of recovery and regime shift reveal.

Stated as a set of postulates, definitions, explicit boundaries, and testable hypotheses, STATE is offered as a foundation to be tested, refined, or rejected by empirical work, rather than as a completed description of autonomic physiology. Static summaries describe where the autonomic system rests; STATE asks what it does when it is pushed.

## Appendix A Proofs and exact results

### A.1 Proof of Proposition 1

*Proof*. (i) Let *Q* be an orthogonal matrix whose first row is *c*^⊤^/‖*c*‖. Because *F* = −*λI* commutes with *Q* and *QW* is again a standard Brownian motion, *X* = *QA* solves *dX* = −*λX dt* + *σ dB*, whose two components are independent Ornstein–Uhlenbeck processes. The output *y* = ‖*c*‖*X*_1_ + *ε* involves only *X*_1_, so *X*_2_ is independent of *y* and its posterior equals its prior. (ii, iii) By the Kalman rank condition, (*F*, *c*^⊤^) is observable if and only if det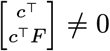; for the stated matrices this determinant equals *c*_*S*_*c*_*P*_ (*λ*_*S*_ − *λ*_*P*_) and *c*_*P*_ [*c*_*S*_(*λ* − *µ*) + *c*_*P*_ *κ*], respectively. ▫

### A.2 Exact properties of the bistable system

The sympathetic coordinate of System B does not depend on *P*_*t*_, so it is an autonomous one-dimensional diffusion for which exact results are available. With *S*_*stress*_ = 2*S*_*saddle*_ =: 2*a*, its drift is −*U*^′^(*S*) with the symmetric double-well potential 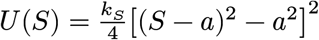.

#### Proposition 2

(exact properties of System B)

Consider *dS*_*t*_ = −*U*^′^(*S*_*t*_) *dt* + *σ dW*_*t*_ with *U* as above, and let *λ* = 2*k*_*S*_*a*^2^ be the rate of System A.

i. The density *π*(*S*) *∝* exp(−2*U*(*S*)/*σ*^2^) is the unique stationary density and is symmetric about *S* = *a*; hence the stationary probability of each basin is 1/2 for every *σ* > 0.
ii. Var_*π*_(*S*) → *a*^2^ as *σ* → 0, whereas the stationary variance of System A is *σ*^2^/(2*λ*) → 0.
iii. The mean first-passage time from the resting attractor to the saddle is

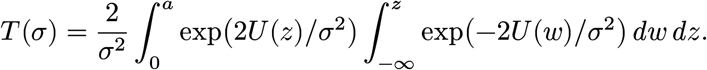
iv. Under a constant forcing *u* > 0 the resting equilibrium exists if and only if 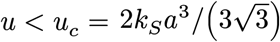, whereas System A requires *u* = *λa* = 2*k*_*S*_*a*^3^ to be displaced to *S* = *a*; the ratio is exactly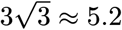.

*Proof*. (i) *U* is coercive and even about *a*, so exp(−2*U*/*σ*^2^) is integrable and symmetric, and for a gradient diffusion with additive noise it is the unique stationary density (Gardiner 2004). (ii) As *σ* → 0, *π* concentrates on the minima {0, 2*a*} with equal weights, so its variance tends to *a*^2^. (iii) This is the standard first-passage formula for a one-dimensional diffusion with a reflecting boundary at −∞ (Gardiner 2004); see also (Hänggi, Talkner, and Borkovec 1990). (iv) The equilibria satisfy *k*_*S*_ *S*(*S* − *a*)(*S* − 2*a*) = *u*; the left-hand side has a local maximum on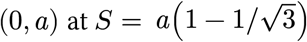, equal to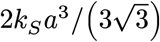, and the resting equilibrium disappears when *u* exceeds it. ▫

For the parameters used here, the exact values are Var_*π*_(*S*) = 0.313 against *σ*^2^/(2*λ*) = 0.0183 for System A (a 17-fold difference), *T* = 217 s at *σ* = 0.10 and *T* = 23.1 s at *σ* = 0.20, and *u*_*c*_ = 0.0316. The pulse used (*u*_*S*_ = 0.50) is therefore 15.8 *u*_*c*_. A Monte Carlo estimate of the first-passage time (3000 replicates, initial state *S* = 0.02) agreed with the exact value within 6%: 47.6 ± 0.8 s against 47.2 s at *σ* = 0.15, and 24.0 ± 0.4 s against 22.8 s at *σ* = 0.20, the small excess being expected from the discrete-time monitoring of the crossing. Proposition 2 has three consequences for any illustration of this kind. First, baseline equivalence is conditional on the window: it can hold only while the baseline is short relative to *T* (*σ*), which is the case at *σ* = 0.10 (25 s against 217 s) but not at *σ* = 0.20. Second, the bistable system is dislodged by forcing 5.2 times smaller than the forcing that displaces its linear counterpart to the same location, so a perturbation of fixed amplitude may saturate the response. Third, the tolerance region must remain inside the resting basin (*δ* < 180 ms), and the saddle (201 ms) and the stress attractor (402 ms) are the natural distance scales against which *δ* must be read.

## Appendix B Simulation specification and robustness

### B.1 Simulation design

Both systems share the parasympathetic equation and the perturbation, and differ in the sympathetic drift *f*_*S*_:

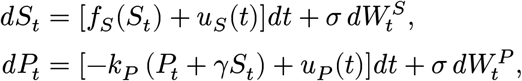

with 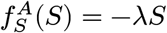 for the monostable System A and 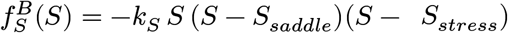 for the bistable System B, where *λ* = *k*_*S*_*S*_*saddle*_*S*_*stress*_. Thus System A is exactly the linearization of System B at rest: both have the same Jacobian at the origin, the same coupling *γ*, and the same noise. We set *k*_*S*_ = 0.38, *S*_*saddle*_ = 0.6, *S*_*stress*_ = 1.2 (so that *λ* = 0.274), *k*_*P*_ = 0.48, *γ* = 0.50, and *σ* = 0.10 in dimensionless units, converted to physiological units by a factor of 300 ms. The deterministic skeleton of System B has a stable resting attractor *ℛ*_*rest*_ = (0, 0), a saddle at (0.6, −0.3) (that is, (180, −90) ms), and a stable stress attractor *ℛ*_*stress*_ = (1.2, −0.6) (that is, (360, −180) ms); the eigenvalues of its Jacobian are {−0.274, −0.48} at both attractors and {+0.137, −0.48} at the saddle. The drift is not a gradient field (*∂f*_*P*_ /*∂S* − *∂f*_*S*_/*∂P* = −*k*_*P*_ *γ* ≠ 0), so no scalar potential exists for the two-dimensional system, consistent with the definition of the landscape in Section 2 the separatrix is the line *S* = *S*_*saddle*_.

At *t* = 25 s an identical transient pulse (*u*_*S*_ = +0.50, *u*_*P*_ = −0.20, that is, *u*_*P*_ = −0.4 *u*_*S*_) was applied to both systems for 20 s, simulating an acute orthostatic or physical stressor. Systems were integrated by the Euler–Maruyama scheme over 100 s (Δ*t* = 0.02 s) from the same initial state.

Displacement was the Euclidean distance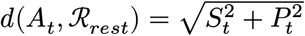. Baseline variability was the standard deviation of *d* over *t* < 25 s; *D*_*max*_ and *T*_*rec*_(*δ, τ*) followed Equation 3 and Equation 4 on a 0.5-s moving average of *d*, with *τ* = 5 s; and *P*_*return*_(*H, τ*) and *P*_*transition*_(*H*) followed Equation 5 and Equation 6 with *H* = 55 s, taking *P*_*transition*_ as the probability 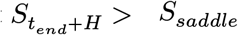 . The tolerance was set by the a priori rule of Section 3.3: *δ* is the 0.999 quantile of the stationary distance of System A, computed exactly from its Gaussian stationary law (the solution of the Lyapunov equation), which gives *δ* = 150.7 ms. This value depends only on the resting regime and lies inside the resting basin, whose half-width is 180 ms. Each system was simulated in 2000 independent replicates; the replicate shown in the state-space panels is the one closest to the ensemble medians of baseline variability, *D*_*max*_, and *T*_*rec*_ (a rule fixed before inspecting trajectories). Exact calculations and simulations are reproducible with the scripts in the repository (see Data availability).

### B.2 Robustness

The contrast depends on the noise regime in the way Appendix A predicts (Figure 3). As *σ* increases from 0.05 to 0.25, the barrier-to-noise ratio falls from 9.8 to 0.4, the proportion of System B replicates in the stress basin at the end of the horizon falls from 100% to 52%, approaching the stationary value of 50%, and the proportion crossing the saddle during the baseline rises from 0% to 83%. Baseline variability then separates the systems increasingly well (AUC rising from 0.53 to 0.70), so baseline equivalence is a property of the low-noise regime and of a short window, and not a general feature of the two systems. The tolerance was kept at *δ* = 150.7 ms at all noise levels. The recovery descriptors separate the systems throughout (AUC for *T*_*rec*_ of at least 0.87).

**Figure 3:**
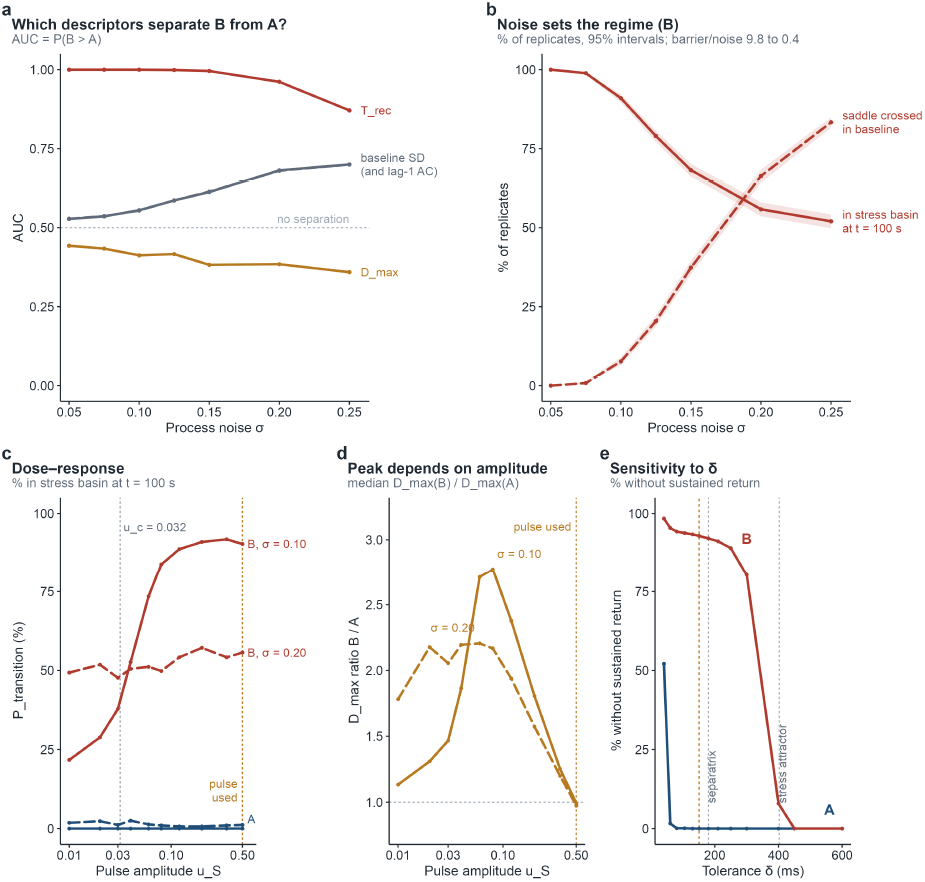
Robustness of the illustration. (a) Separation of A and B (AUC; 0.5 = none) by descriptor across noise levels *σ* (2000 replicates per level). (b) Proportion of B replicates in the stress basin at *t* = 100 s and in which the saddle was crossed during the baseline, with 95% Wilson intervals; the barrier-to-noise ratio Δ*U*/(*σ*^2^/2) falls from 9.8 to 0.4 across the range. (c) Dose– response of the regime shift to the pulse amplitude *u*_*S*_ (*u*_*P*_ = −0.4 *u*_*S*_; 600 replicates per point) in System B (red) and System A (blue) at *σ* = 0.10 (solid) and 0.20 (dashed); dotted lines mark the exact critical forcing *u*_*c*_ and the pulse of Figure 2. (d) Ratio of the median peak displacement of B to that of A across amplitudes. (e) Proportion of replicates without sustained return as a function of *δ* (1000 replicates; *σ* = 0.10; red: B, blue: A); the dashed line is the rule value, and the dotted lines mark the separatrix (180 ms) and the stress attractor (402 ms).

The dose–response confirms this reading and qualifies the comparison of peaks. At *σ* = 0.10, the proportion of System B replicates shifting regime rises from 22% at *u*_*S*_ = 0.01 to 53% at *u*_*S*_ = 0.04 (close to *u*_*c*_ = 0.0316) and plateaus at about 90% for *u*_*S*_ ≥ 0.12; System A almost never shifts regime. At *σ* = 0.20 it is 48–57% irrespective of amplitude, which reflects noise-induced equilibration and not a response to the perturbation. The coincidence of peaks, in contrast, is specific to the largest amplitude: at *σ* = 0.10 the median *D*_*max*_ of System B exceeds that of System A by a factor of 1.31 at *u*_*S*_ = 0.02, 1.86 at 0.04, 2.77 at 0.08, 1.81 at 0.20, and 1.25 at 0.35, and the two become equal only at *u*_*S*_ = 0.5 (ratio 0.99), because once the separatrix is crossed System B moves toward the second attractor, 402 ms from rest, whereas the linear response grows only in proportion to the forcing. Peak displacement is therefore protocol-dependent and cannot by itself identify the regime; the AUC of 0.36–0.44 for *D*_*max*_ across noise levels in Figure 3 applies to the pulse used.

The tolerance also matters: no sustained return was observed in at least 80% of System B replicates for any *δ* ≤ 300 ms, in 7.9% at *δ* = 400 ms, and in none at *δ* ≥ 450 ms, because the stress attractor lies 402 ms from *ℛ*_*rest*_; at *δ* = 40 ms even System A failed to return (52%), because *δ* then lies within its own stationary fluctuations. Finally, reducing the integration step from 0.02 s to 0.005 s changed the median *D*_*max*_ by less than 1% and *P*_*transition*_ by 1.7 percentage points (600 replicates), within Monte Carlo error.

## CRediT authorship contribution statement

Matías Castillo-Aguilar: Conceptualization, Methodology, Investigation, Visualization, Writing – original draft, Writing – review & editing. Diego Mabe-Castro: Writing – original draft, Writing – review & editing. Cristian Núñez-Espinosa: Writing – original draft, Writing – review & editing, Supervision, Funding acquisition.

## Declaration of competing interest

The authors declare no conflicts of interest.

## Funding

This research was funded by ANID Proyecto FONDECYT Regular, grant number 1250474, received by C.N.-E. The funder had no role in the study design, analysis, interpretation of results, writing of the report, or the decision to submit the article for publication.

## Data availability

The code and results presented in this study are openly available in the GitHub repository https://github.com/matcasti/STATE-paper. All exact calculations, simulations, tables, and figures can be regenerated with `R/run-all.R;` the simulation seeds, parameters, and software versions are recorded in `results/.` No empirical data were used.

## Declaration of generative AI and AI-assisted technologies in the manuscript preparation process

During the preparation of this work the authors used Claude (Anthropic) for language editing and to assist with the code and text of the computational illustration. After using this tool, the authors reviewed and edited the content as needed and take full responsibility for the content of the published article.

## Notes

https://github.com/matcasti/STATE-paper

### Competing Interest Statement

The authors have declared no competing interest.

## References

Ao, Ping. 2004. “Potential in Stochastic Differential Equations: Novel Construction.” Journal of Physics A: Mathematical and General 37 (3): L25–30. 10.1088/0305-4470/37/3/L01.

Barabási, Albert-László, Natali Gulbahce, and Joseph Loscalzo. 2011. “Network Medicine: A Network-Based Approach to Human Disease.” Nature Reviews Genetics 12 (1): 56–68. 10.1038/nrg2918.

Bartsch, Ronny P., Kang K. L. Liu, Amir Bashan, and Plamen Ch. Ivanov. 2015. “Network Physiology: How Organ Systems Dynamically Interact.” PLoS ONE 10 (11): e0142143. 10.1371/journal.pone.0142143.

Benarroch, Eduardo E. 1993. “The Central Autonomic Network: Functional Organization, Dysfunction, and Perspective.” Mayo Clinic Proceedings 68 (10): 988–1001. 10.1016/S0025-6196(12)62272-1.

Castillo-Aguilar, Matías, David Medina-Ortiz, Ruby Méndez Muñoz, Diego Mabe-Castro, Noah Beelders, Atenea Uribe-Ojeda, Marcelo A. Navarrete, and Cristian Núñez-Espinosa. 2026. “BAND: A Probabilistic Framework for Modeling Non-Stationary Heart Rate Variability in Rest–Stress– Rest Dynamics.” Technologies 14 (9): 528. 10.3390/technologies14090528.

Cole, Christopher R., Eugene H. Blackstone, Fredric J. Pashkow, Claire E. Snader, and Michael S. Lauer. 1999. “Heart-Rate Recovery Immediately After Exercise as a Predictor of Mortality.” New England Journal of Medicine 341 (18): 1351–57. 10.1056/NEJM199910283411804.

Costa, Madalena, Ary L. Goldberger, and C.-K. Peng. 2002. “Multiscale Entropy Analysis of Complex Physiologic Time Series.” Physical Review Letters 89 (6): 068102. 10.1103/PhysRevLett.89.068102.

Eckberg, Dwain L. 1983. “Human Sinus Arrhythmia as an Index of Vagal Cardiac Outflow.” Journal of Applied Physiology 54 (4): 961–66. 10.1152/jappl.1983.54.4.961.

Gardiner, Crispin W. 2004. Handbook of Stochastic Methods for Physics, Chemistry and the Natural Sciences. 3rd ed. Berlin: Springer.

Gaur, Pooja, Dorota S. Temple, Meghan Hegarty-Craver, Matthew D. Boyce, Jonathan R. Holt, Michael F. Wenger, Edward A. Preble, et al. 2024. “Continuous Monitoring of Heart Rate Variability in Free-Living Conditions Using Wearable Sensors: Exploratory Observational Study.” JMIR Formative Research 8: e53977. 10.2196/53977.

Ghazzal, Bahjat Z., Ibrahim Hasan, Nebras Hasan, Ziyad Ghazzal, and Marwan M. Refaat. 2026. “Interpreting Heart Rate Variability in Clinical Context: A Framework for Understanding a Multidimensional Autonomic Biomarker.” Frontiers in Cardiovascular Medicine 13: 1949249. 10.3389/fcvm.2026.1949249.

Gijzel, Sanne M. W., Ingrid A. van de Leemput, Marten Scheffer, Mattia Roppolo, Marcel G. M. Olde Rikkert, and René J. F. Melis. 2017. “Dynamical Resilience Indicators in Time Series of Self-Rated Health Correspond to Frailty Levels in Older Adults.” The Journals of Gerontology: Series A 72 (7): 991–96. 10.1093/gerona/glx065.

Hänggi, Peter, Peter Talkner, and Michal Borkovec. 1990. “Reaction-Rate Theory: Fifty Years After Kramers.” Reviews of Modern Physics 62 (2): 251–341. 10.1103/RevModPhys.62.251.

Holling, C. S. 1973. “Resilience and Stability of Ecological Systems.” Annual Review of Ecology and Systematics 4: 1–23. 10.1146/annurev.es.04.110173.000245.

Jandackova, Vera K., Shaun Scholes, Annie Britton, and Andrew Steptoe. 2016. “Are Changes in Heart Rate Variability in Middle-Aged and Older People Normative or Caused by Pathological Conditions? Findings from a Large Population-Based Longitudinal Cohort Study.” Journal of the American Heart Association 5 (2): e002365. 10.1161/JAHA.115.002365.

Kemp, Andrew H., Daniel S. Quintana, Marcus A. Gray, Kim L. Felmingham, Kerri Brown, and Justine M. Gatt. 2010. “Impact of Depression and Antidepressant Treatment on Heart Rate Variability: A Review and Meta-Analysis.” Biological Psychiatry 67 (11): 1067–74. 10.1016/j.biopsych.2009.12.012.

Kuusela, Tom A., Tony Shepherd, and Jarmo Hietarinta. 2003. “Stochastic Model for Heart-Rate Fluctuations.” Physical Review E 67: 061904. 10.1103/PhysRevE.67.061904.

Leemput, Ingrid A. van de Marieke Wichers, Angélique O. J. Cramer, Denny Borsboom, Francis Tuerlinckx, Peter Kuppens, Egbert H. van Nes, et al. 2014. “Critical Slowing down as Early Warning for the Onset and Termination of Depression.” Proceedings of the National Academy of Sciences 111 (1): 87–92. 10.1073/pnas.1312114110.

Mackey, Michael C., and Leon Glass. 1977. “Oscillation and Chaos in Physiological Control Systems.” Science 197 (4300): 287–89. 10.1126/science.267326.

McEwen, Bruce S., and Eliot Stellar. 1993. “Stress and the Individual: Mechanisms Leading to Disease.” Archives of Internal Medicine 153 (18): 2093–2101. 10.1001/archinte.1993.00410180039004.

Olde Rikkert, Marcel G. M., Vasilis Dakos, Timothy G. Buchman, Rudolf de Boer, Leon Glass, Angélique O. J. Cramer, Simon Levin, et al. 2016. “Slowing down of Recovery as Generic Risk Marker for Acute Severity Transitions in Chronic Diseases.” Critical Care Medicine 44 (3): 601–6. 10.1097/CCM.0000000000001450.

Onnela, Jukka-Pekka, and Scott L. Rauch. 2016. “Harnessing Smartphone-Based Digital Phenotyping to Enhance Behavioral and Mental Health.” Neuropsychopharmacology 41 (7): 1691–96. 10.1038/npp.2016.7.

Percha, Bethany, Edward B. Baskerville, Matthew Johnson, Joel T. Dudley, and Noah Zimmerman. 2019. “Designing Robust n-of-1 Studies for Precision Medicine: Simulation Study and Design Recommendations.” Journal of Medical Internet Research 21 (4): e12641. 10.2196/12641.

Qiu, Shanhu, Xue Cai, Zilin Sun, Ling Li, Martina Zuegel, Juergen M. Steinacker, and Uwe Schumann. 2017. “Heart Rate Recovery and Risk of Cardiovascular Events and All-Cause Mortality: A Meta-Analysis of Prospective Cohort Studies.” Journal of the American Heart Association 6 (5): e005505. 10.1161/JAHA.117.005505.

Rector, Jerrald L., Sanne M. W. Gijzel, Ingrid A. van de Leemput, Fokke B. van Meulen, Marcel G. M. Olde Rikkert, and René J. F. Melis. 2021. “Dynamical Indicators of Resilience from Physiological Time Series in Geriatric Inpatients: Lessons Learned.” Experimental Gerontology 149: 111341. 10.1016/j.exger.2021.111341.

Rosas, Fernando E., Diego Candia-Rivera, Andrea I. Luppi, Yike Guo, and Pedro A. M. Mediano. 2024. “Bayesian at Heart: Towards Autonomic Outflow Estimation via Generative State-Space Modelling of Heart Rate Dynamics.” Computers in Biology and Medicine 170: 107857. 10.1016/j.compbiomed.2023.107857.

Scheffer, Marten, Jordi Bascompte, William A. Brock, Victor Brovkin, Stephen R. Carpenter, Vasilis Dakos, Hermann Held, Egbert H. van Nes, Max Rietkerk, and George Sugihara. 2009. “Early-Warning Signals for Critical Transitions.” Nature 461 (7260): 53–59. 10.1038/nature08227.

Shaffer, Fred, and J. P. Ginsberg. 2017. “An Overview of Heart Rate Variability Metrics and Norms.” Frontiers in Public Health 5: 258. 10.3389/fpubh.2017.00258.

Sundas, Amina, Ivan Contreras, Judith Navarro-Otano, Júlia Soler, Aleix Beneyto, and Josep Vehi. 2025. “Heart Rate Variability over the Decades: A Scoping Review.” PeerJ 13: e19347. 10.7717/peerj.19347.

Task Force of the European Society of Cardiology and the North American Society of Pacing and Electrophysiology. 1996. “Heart Rate Variability: Standards of Measurement, Physiological Interpretation, and Clinical Use.” Circulation 93 (5): 1043–65. 10.1161/01.CIR.93.5.1043.

Thayer, Julian F., and Richard D. Lane. 2000. “A Model of Neurovisceral Integration in Emotion Regulation and Dysregulation.” Journal of Affective Disorders 61 (3): 201–16. 10.1016/S0165-0327(00)00338-4.

Wang, Jin, Li Xu, and Erkang Wang. 2008. “Potential Landscape and Flux Framework of Nonequilibrium Networks: Robustness, Dissipation, and Coherence of Biochemical Oscillations.” Proceedings of the National Academy of Sciences 105. 10.1073/pnas.0800579105.

Wieland, Franz-Georg, Adrian L. Hauber, Marcus Rosenblatt, Christian Tönsing, and Jens Timmer. 2021. “On Structural and Practical Identifiability.” Current Opinion in Systems Biology 25: 60–69. 10.1016/j.coisb.2021.03.005.

Wild, Christopher P. 2005. “Complementing the Genome with an “Exposome”: The Outstanding Challenge of Environmental Exposure Measurement in Molecular Epidemiology.” Cancer Epidemi/ ology, Biomarkers & Prevention 14 (8): 1847–50. 10.1158/1055-9965.EPI-05-0456.

